# Kinome Profiling of Primary Endometrial Tumors Using Multiplexed Inhibitor Beads and Mass Spectrometry Identifies SRPK1 As Candidate Therapeutic Target

**DOI:** 10.1101/2020.03.03.970251

**Authors:** Katherine J. Johnson, Vikas Kumar, Alison M. Kurimchak, Nishi Srivastava, Suraj Peri, Kathy Q. Cai, Gina M. Mantia-Smaldone, James S. Duncan

## Abstract

Protein kinases (collectively, termed the kinome) represent one of the most tractable drug targets in the pursuit of new and effective cancer treatments. However, less than 20% of the kinome is currently being explored as primary targets for cancer therapy, leaving the majority of the kinome untargeted for drug therapy. Chemical proteomics approaches such as Multiplexed Inhibitor Beads and Mass Spectrometry (MIB-MS) have been developed that measure the abundance of a significant portion of the kinome, providing a strategy to interrogate kinome landscapes and dynamics. Kinome profiling of cancer cell lines using MIB-MS has been extensively characterized, however, application of this method to measure tissue kinome(s) has not been thoroughly explored. Here, we present a quantitative proteomics workflow specifically designed for kinome profiling of tissues that pairs MIB-MS with a newly designed super-SILAC kinome standard. Using this workflow, we mapped the kinome landscape of endometrial carcinoma (EC) tumors and normal endometrial (NE) tissues and identified several kinases overexpressed in EC tumors, including Serine/Arginine-Rich Splicing Factor kinase, (SRPK1). Immunohistochemical (IHC) analysis of EC tumor TMAs confirmed MIB-MS findings and showed SRPK1 protein levels were highly expressed in endometrioid and uterine serous cancer (USC) histological subtypes. Querying large-scale genomics studies of EC tumors revealed high expression of SRPK1 correlated with poor survival. Inhibition of SRPK1 in USC cells altered mRNA splicing, downregulating several oncogenes including MYC and Survivin resulting in apoptosis. Taken together, we present a SILAC-based MIB-MS kinome profiling platform for measuring kinase abundance in tumor tissues, and demonstrate its application to identify SRPK1 as a plausible kinase drug target for the treatment of EC.

## INTRODUCTION

Protein kinases are a family of ~535 enzymes that, collectively, are termed the kinome (*1*). Uncontrolled protein kinase activity has been linked to the development of nearly 25% of all cancers; consequently, protein kinases represent one of the most promising avenues for cancer therapy (*2, 3*). Indeed, >30 kinase-specific inhibitors are currently approved for therapy of various cancer types, with more than 150 kinase inhibitors in Phase 1-3 clinical trials across all diseases (*4*). However, most of these kinase-specific inhibitors target a relatively small fraction of the human kinome with only about 20% avidly being explored as primary targets for drug therapy (*5, 6*). Thus, the majority of the kinome remains untargeted for cancer therapy and about 50% of the kinome is largely uncharacterized with respect to the function and role of these kinases in cancer, representing the understudied or ‘dark’ cancer kinome (*6–8*). Notably, several CRISPR/cas9 and/or RNAi loss-of-function studies have shown that many dark kinases are essential for cancer cell viability highlighting the therapeutic potential of the dark kinome for the treatment of cancer (*9, 10*).

Identifying kinase-signaling networks that are essential for tumor growth and resistance requires a detailed knowledge of global kinome activity (sum of all 518 human kinases), not simply measuring one or a few kinases in a pathway. To accomplish this, quantitative proteomics strategies including Kinobeads, Kinativ and Multiplexed Inhibitor Beads (MIBs) have been developed that are capable of detecting dynamic changes globally across the kinome at baseline or following treatment with a targeted inhibitor (*11–13*). These chemical proteomics techniques couple kinase affinity capture with quantitative mass spectrometry, providing a systems biology platform to profile global kinome signaling at the proteomic level. Several reports have applied kinome enrichment strategies to define kinase signatures of cancer cell lines (*14*), however, a robust quantitative kinome enrichment strategy for measuring kinase abundance across cohorts of tissues, (i.e., primary tumors) has not been systematically explored. As the usage of patient-derived xenografts (PDX) models (*15*) for drug studies increases, and frozen tissue samples from clinical trials become more frequently available for analysis, a tumor kinome profiling strategy specifically designed for tissue analysis will be of particular interest.

Here, we present an optimized Quantitative-Multiplexed Inhibitor Beads (Q-MIBs) workflow for profiling the kinome in tissues that incorporates a newly-designed super-SILAC kinome reference standard permitting robust quantitation of ~70% of the human kinome. Following quality control testing, we applied Q-MIBs to identify candidate kinase targets in endometrial carcinoma (EC), which is the most common gynecologic malignancy in the United States with limited effective targeted therapies (*16*). Using Q-MIBs, we profiled the kinome of primary patient endometrial carcinoma (EC) tumors and normal endometrial (NE) tissues and identified overexpression of a network of kinases that included Serine/Arginine-Rich Splicing Factor kinase, (SRPK1). Inhibition of SRPK1 in a recognized USC cell line caused significant alternative splicing downregulating several established EC oncogenes including MYC, and Survivin inducing apoptosis. These studies demonstrated the applicability of Q-MIBs to measure kinase abundance in tumor tissues, and nominated SRPK1 as a plausible therapeutic target for the treatment of ECs.

## EXPERIMENTAL PROCEDURES

### Experimental Design, Data Analysis and Statistical Rationale

For proteomic measurement of kinase abundance in tissues, five milligrams of each primary tissue were run on a single MIB-column, followed by LC-MS/MS analysis in technical replicates. An equal amount of the s-SILAC kinome standard reference (SKS) was spiked into non-SILAC labeled tissue samples, endogenous kinases purified by MIB-resins, kinases eluted, digested, and peptides analyzed by LC-MS/MS. Kinase protein levels in tissues were quantitated by comparing SILAC-labeled peptides from SKS with non-labeled peptides from tissues using MaxQuant. To increase the statistical power, 20 distinct primary endometrial tumors and 20 distinct normal endometrial tissues were measured by Q-MIBs. Notably, 6 of the normal endometrial tissues and 2 of the endometrial tumors yielded insufficient kinases measurement (<180 measured) and were excluded from the comparative analysis. For MIB-MS data analysis, MaxQuant normalized ratios were imported into Perseus software (1.6.2.3) for quantitation. MIB-MS profiles were processed in Perseus software the following manner: normalized MIB-MS s-SILAC ratios were transformed 1/(x) to generate light / heavy ratios, followed by log2/(1/x) transformed. Columns were then filtered based on a valid value of 150, MIB-MS technical replicates averaged, and rows filtered for minimum valid kinases measured (n=>3 kinases). Imputation of missing values was performed as previously described (*17*), where in the s-SILAC data, a width of 0.3 and the downshift of 0.5, was employed. Principal component analysis (PC1 vs PC2, PC2 vs PC3 and PC1 vs PC3) was then performed to visualize kinome profiles amongst samples. Hierarchical clustering (Euclidean) of s-SILAC ratios was performed and column clusters annotated selecting 4 clusters at a distance threshold of 27.87. Difference in kinase abundance amongst endometrial tumors (n=17) and normal endometrial tissues (n=14) was determined using a two-sample Student’s t-test with the following parameters, (S0 0.1, and Side, Both) using p-value < 0.05 using Perseus software. Analysis of SRPK1 mRNA alterations in endometrial tumors (n=232) from TCGA studies (*17*) association with survival was performed using Fisher’s two-sided p < 0.05. Immunohistochemistry analysis of overexpression of SRPK1 in endometrioid (n=18) and serous (39) endometrial tumors relative to normal endometrial tissues (n=12) was determined by two-sample Student’s t-test P < 0.01. For splicing analysis following SPHINX3.1 treatment, RNA-seq reads for both treatment and controls were aligned to the human genome (Hg38) using TopHat (*18*). Detection of splicing changes between treatment and controls was assessed using MATS algorithm, using the aligned BAM files with default parameters (*19*). The spliced events were filtered using False Discovery Rate by Benjamini-Hochberg method (FDR < 0.01). Genes significantly altered by SPHINX31 treatment in SPEC-2 cells were determined by two-sample Student’s t-test P < 0.001 comparing DMSO vs SPHINX31-treatment and analyzed using g:Profiler and Database for Annotation, Visualization and Integrated Discovery (DAVID) v6.8 to determine pathway enrichments (BH P<05).

### Cell Lines

The SPEC-2 cell line was generously provided by Chunxiao Zhou, University of North Carolina. All cell lines were verified by IDEXX laboratories and were verified Mycoplasma-negative (1/7/20) using the Hoechst DNA stain method. SPEC-2, SW48, MOLT4, UACC257, ACHN, COLO205, SF295, SKOV3, OVCAR5 and PC-3 cells were maintained in RPMI-1640 supplemented with 10% FBS, 100 U/ml Penicillin-Streptomycin and 2mM GlutaMAX. Cell lines used for the s-SILAC reference sample (MOLT4, UACC257, ACHN, COLO205, SF295, SKOV3, OVCAR5 and PC-3) were grown for seven doublings in arginine- and lysine-depleted media supplemented with heavy isotope labeled [^13^C_6_, ^15^N_4_]arginine (Arg10) (84 mg/L) and [^13^C_6_]lysine (Lys8) (48 mg/L) (Sigma), and unlabeled leucine (50 mg/L) (ThermoFisher Scientific) as described previously (*20*). All cells were kept at 37°C in a 5% CO_2_ incubator.

### Patient Samples

All patient samples used in this study were obtained from the Fox Chase Cancer Center (FCCC) Biosample Repository Facility (BRF). The FCCC BRF maintains a longstanding Institutional Review Board-approved protocol for collection, banking and distribution of de-identified biospecimens and associated clinical data. Consent and authorization for the use of de-identified specimens and associated clinical data for unrestricted research was obtained from all BRF participants prior to specimen collection.

### Frozen Tumor Samples

Endometrial tumors and matched normal adjacent endometrial tissues were obtained from the FCCC BRF for this IRB-approved research (16-9031 and 14-809). Tumor tissue was snap frozen at the time of collection and stored at −80°C in the FCCC BRF. Tumor histology and cellularity was confirmed by the FCCC BRF pathologist at the time of banking.

### Tumor Xenograft Studies

SW48 colorectal cancer cells were implanted in Nude mice and grown as tumor xenografts. Tumors were harvested when tumors were approximately 200 mm^3^ and immediately snap frozen in liquid nitrogen. All studies were conducted after review by Fox Chase Cancer Intuitional Animal Care and Use Committee (IACUC, # 16-16).

### Compounds

SPHINX31 was purchased from Axon Medchem (#2714). For the compounds utilized in MIB synthesis, Purvalanol B was purchased from Abcam. PP58 and VI16832 were custom synthesized according to previously described methods by The Center for Combinatorial Chemistry and Drug Discovery, Jilin University, P.R. China (*21*). CTx-0294885 was purchased from MedKoo Biosciences, Inc (406457). Conjugation of inhibitors to beads was performed by carbodiimide coupling to ECH Sepharose 4B (CTx-0294885, VI16832 and PP58) or EAH Sepharose 4B (purvalanol B) (GE Healthcare).

### Immunoblotting

Samples were harvested in MIB lysis buffer, subjected to SDS-PAGE chromatography and transferred to PVDF membranes before western blotting with primary antibodies. Primary antibodies Cyclin D1 (#2978), Survivn (#2808), GAPDH (#2118), MYC (#5065), PARP (#9542) were purchased from Cell Signaling Technologies. Phospho-SRSF (#MABE50) was purchased from EMD Millipore, and SRPK1 (#ab90527) was purchased from Abcam. Secondary HRP-anti-rabbit and HRP-anti-mouse were obtained from ThermoFisher Scientific. SuperSignal West Pico and Femto Chemiluminescent Substrates (Thermo Scientific) were used to visualize blots.

### Endometrial Tissues and Tumor Specimens for IHC

Cancerous and benign endometrial specimens were obtained from patients who underwent surgical resection at Fox Chase Cancer Center. Two endometrial tumor tissue microarrays (TMAs) containing a total of 57 EC patient tumors, (39 USC and 18 endometrioid) and 12 normal endometrial tissues in duplicate were provided by the Biosample Repository of Fox Chase Cancer Center (Data file S5). All these samples were used with informed patient consent, and the study was approved by the Institutional Review Boards of Fox Chase Cancer Center. Typically, tissue specimens were fixed in 10% buffered formalin, embedded in paraffin, sectioned, and stained with hematoxylin and eosin. All tumors were histologically classified according to the World Health Organization (WHO) classification, and the surgical stages were determined according to the classification of International Federation of Gynecology and Obstetrics (FIGO).

### Immunohistochemistry (IHC) and IHC Evaluation

Immunohistochemical staining was carried out according to standard methods by the FCCC Histopathology Facility. Briefly, 5µm formalin-fixed, paraffin-embedded TMA sections were deparaffinized and hydrated. Sections were then subjected to heat-induced epitope retrieval with 0.01 M citrate buffer (pH 6.0). Endogenous peroxidases were quenched by the immersion of slides in 3% hydrogen peroxide solution. The sections were incubated overnight with primary antibodies to SRPK1 (Rabbit, 1:200, HPA016431, Sigma-Aldrich) at 4 °C in a humidified slide chamber. Immunodetection was performed using the Dako Envision+ polymer system and immunostaining was visualized with the chromogen 3, 3’-diaminobenzidine. The sections were then washed, counterstained with hematoxylin, dehydrated with ethanol series, cleared in xylene, and mounted. Known positive cancer tissues were used as positive controls. As a negative control, the primary antibody was replaced with normal rabbit IgG to confirm absence of specific staining. SRPK1 stained TMA slides were scanned with Leica AperioScanScope CS2. Scanned images were then viewed and captured with Aperio’s image viewer software (ImageScope, version 12.4). Immunoreactivity of SRPK1 protein was evaluated by a pathologist and H score was generated. The H score consists of the product of the intensity of cytoplasmic staining (0-3+) and the percentage of cells with positive cytoplasmic staining. The range of the H score was 0-300 (0% cells positive to 100% cells 3+positive).

### MIBs Preparation and Chromatography

Experiments using MIB/MS were performed as previously described (*13*). Briefly, cells or tumors were lysed on ice in buffer containing 50 mM HEPES (pH 7.5), 0.5% Triton X-100, 150 mM NaCl, 1 mM EDTA, 1 mM EGTA, 10 mM sodium fluoride, 2.5 mM sodium orthovanadate, 1X protease inhibitor cocktail (Roche), and 1% each of phosphatase inhibitor cocktails 2 and 3 (Sigma). Particulate was removed by centrifugation of lysates at 21,000 g for 15 minutes at 4°C and filtration through 0.45 µm syringe filters. Protein concentrations were determined by BCA analysis (Thermo Scientific). To compare kinome signatures of human tissues, we designed a super-SILAC method consisting of a cocktail of cancer cell lines that encompasses the activity of the “cancer kinome” to be used as a control for individual samples. Five cancer cell lines were selected from the NCI-60 panel (UACC257, MOLT4, COLO205, ACHN and PC3) that differed in their origin, gene expression patterns and mutation status. The cancer cell lines were labeled using SILAC and mixed equally providing a diverse SILAC ([^13^C_6_, ^15^N_4_] arginine (Arg 10) and [^13^C_6_] lysine (Lys 8)) heavy reference standard (s-SILAC) to be used repeatedly in kinome profiling assays. An equal amount of the s-SILAC reference (5 mg) lysate was added to our non-labeled (5 mg) sample (cell, or tumor tissue) and analyzed on MIB-beads. Endogenous kinases were isolated by flowing lysates over kinase inhibitor-conjugated Sepharose beads (purvalanol B, VI16832, PP58 and CTx-0294885 beads) in 10 ml gravity-flow columns. After 2×10 ml column washes in high-salt buffer and 1×10 ml wash in low-salt buffer (containing 50 mM HEPES (pH 7.5), 0.5% Triton X-100, 1 mM EDTA, 1 mM EGTA, and 10 mM sodium fluoride, and 1M NaCl or 150 mM NaCl, respectively), retained kinases were eluted from the column by boiling in 2 x 500 µl of 0.5% SDS, 0.1 M TrisHCl (pH 6.8), and 1% 2-mercaptoethanol. Eluted peptides were reduced by incubation with 5 mM DTT at 65°C for 25 minutes, alkylated with 20 mM iodoacetamide at room temperature for 30 minutes in the dark, and alkylation was quenched with DTT for 10 minutes. Samples were concentrated to approximately 100 µl with Millipore 10kD cutoff spin concentrators. Detergent was removed by chloroform/methanol extraction, and the protein pellet was resuspended in 50 mM ammonium bicarbonate and digested with sequencing-grade modified trypsin (Promega) overnight at 37°C. Peptides were cleaned with PepClean C18 spin columns (ThermoFisher Scientific) dried in a speed-vac, resuspended in 50 μl 0.1% formic acid, and extracted with ethyl acetate (10:1 ethyl acetate:H_2_O). Briefly, 1 mL ethyl acetate was added to each sample, vortexed and centrifuged at max speed for 5 minutes, then removed. This process is repeated 4 more times. After removal of ethyl acetate following the 5th centrifugation, samples were placed at 60°C for 10 minutes to evaporate residual ethyl acetate. The peptides were dried in a speed vac, and subsequent LC-/MS/MS analysis was performed.

### Nano-LC-MS/MS

Proteolytic peptides were resuspended in 0.1% formic acid and separated with a Thermo Scientific RSLC Ultimate 3000 on a Thermo Scientific Easy-Spray C18 PepMap 75µm x 50cm C-18 2 μm column with a 240 min gradient of 4-25% acetonitrile with 0.1% formic acid at 300 nL/min at 50°C. Eluted peptides were analyzed by a Thermo Scientific Q Exactive plus mass spectrometer utilizing a top 15 methodology in which the 15 most intense peptide precursor ions were subjected to fragmentation. The AGC for MS1 was set to 3×10^6^ with a max injection time of 120 ms, the AGC for MS2 ions was set to 1×10^5^ with a max injection time of 150 ms, and the dynamic exclusion was set to 90 s.

### Data Processing

Raw data analysis of SILAC experiments was performed using Maxquant software 1.6.1.0 and searched using andromeda 1.5.6.0 against the swiss-prot human protein database (downloaded on July 26, 2018). The search was set up for full tryptic peptides with a maximum of two missed cleavage sites. All settings were default and searched using acetylation of protein N-terminus and oxidized methionine as variable modifications. Carbamidomethylation of cysteine was set as fixed modification. The precursor mass tolerance threshold was set at 10 ppm and maximum fragment mass error was 0.02 Da. SILAC quantification was performed using MaxQuant by choosing multiplicity as 2 in group-specific parameters and Arg10 and Lys8 as heavy labels. The match between runs was employed and the significance threshold of the ion score was calculated based on a false discovery rate of < 1%.

### RNA Sequencing and Alternative Splicing Analysis

RNA library preparations and sequencing reactions were conducted at GENEWIZ, LLC. (South Plainfield, NJ, USA). RNA samples received were quantified using Qubit 2.0 Fluorometer (Life Technologies, Carlsbad, CA, USA) and RNA integrity was checked using Agilent TapeStation 4200 (Agilent Technologies, Palo Alto, CA, USA). RNA sequencing libraries were prepared using the NEBNext Ultra RNA Library Prep Kit for Illumina following manufacturer’s instructions (NEB, Ipswich, MA, USA). Briefly, mRNAs were first enriched with Oligo(dT) beads. Enriched mRNAs were fragmented for 15 minutes at 94 °C. First strand and second strand cDNAs were subsequently synthesized. cDNA fragments were end repaired and adenylated at 3’ends, and universal adapters were ligated to cDNA fragments, followed by index addition and library enrichment by limited-cycle PCR. The sequencing libraries were validated on the Agilent TapeStation (Agilent Technologies, Palo Alto, CA, USA), and quantified by using Qubit 2.0 Fluorometer (Invitrogen, Carlsbad, CA) as well as by quantitative PCR (KAPA Biosystems, Wilmington, MA, USA). The sequencing libraries were clustered on 1 lane of a flowcell. After clustering, the flowcell was loaded on the Illumina HiSeq instrument (4000 or equivalent) according to manufacturer’s instructions. The samples were sequenced using a 2×150bp Paired End (PE) configuration. Image analysis and base calling were conducted by the HiSeq Control Software (HCS). Raw sequence data (.bcl files) generated from Illumina HiSeq was converted into fastq files and de-multiplexed using Illumina’s bcl2fastq 2.17 software. One mismatch was allowed for index sequence identification. RNA sequencing reads for both treatment and controls were aligned to the human genome (hg38) using TopHat. (19). Alternative splicing analysis was performed as described previously (*22*). Briefly, for detection of splicing changes between treatment and controls, the MATS algorithm was implemented using the aligned BAM files with default parameters (*19*). The spliced events were filtered using False Discovery Rate by Benjamini-Hochberg method (FDR < 0.01). Furthermore, the events were sorted based on difference between average of inclusion level for treated and control samples. Each splicing change was visualized using the IGV program (Integrative Genomics Viewer). Enrichment analysis for Gene Ontology (GO) terms was assessed using the GOstats program (*23*).

### Bioinformatics Analyses of TCGA Datasets

Analysis of SRPK1 CNA and mRNA alterations in EC from TCGA studies was performed at the Biostatistics and Bioinformatics Facility, FCCC. SRPK1 expression is associated with survival (Fisher’s two-sided p < 0.05). Overall survival was based on vital status and “days to death” from initial pathologic diagnosis. Individuals who were still alive at the time of the last follow-up were censored. Survival curves were compared with log-rank tests, and these calculations were done using the R ‘survival’ package (Therneau, T. M. & Grambsch, P. M. Modeling Survival Data: Extending the Cox Model (Springer-Verlag, 2010). Survival data was obtained from TCGA data repository.

## RESULTS

### Designing a Super-SILAC Kinome Reference Standard for Measuring MIB-enriched Kinase Abundance in Tissues

Kinome profiling strategies have been widely utilized in cancer cell line models, however, application of kinase enrichment strategies to measure kinase abundance in tumor tissues has not been extensively characterized. Here, to quantitate the levels of protein kinases in tissue samples, we designed a super-SILAC (*24*) kinome standard (SKS) that can be paired with Multiplexed Inhibitor Beads and Mass Spectrometry (*12*) to measure kinase abundance across samples, which we’ve termed Quantitative-Multiplexed Inhibitor Beads (Q-MIBs) (Fig. 1). An equal amount of the SKS is spiked into any non-SILAC labeled sample (snap frozen tissues), endogenous kinases purified by MIBs, kinases eluted, digested, and peptides analyzed by LC-MS/MS. Kinase protein levels in tissues are then quantitated by comparing SILAC-labeled peptides from SKS with non-labeled peptides from tissues using MaxQuant Software (*25*). Collectively, the Q-MIBs workflow generates kinase signatures that can be immediately actionable through small molecules, as well as identifying perturbations of understudied kinases, which could represent new drug targets for cancer therapy.

**Fig. 1.**
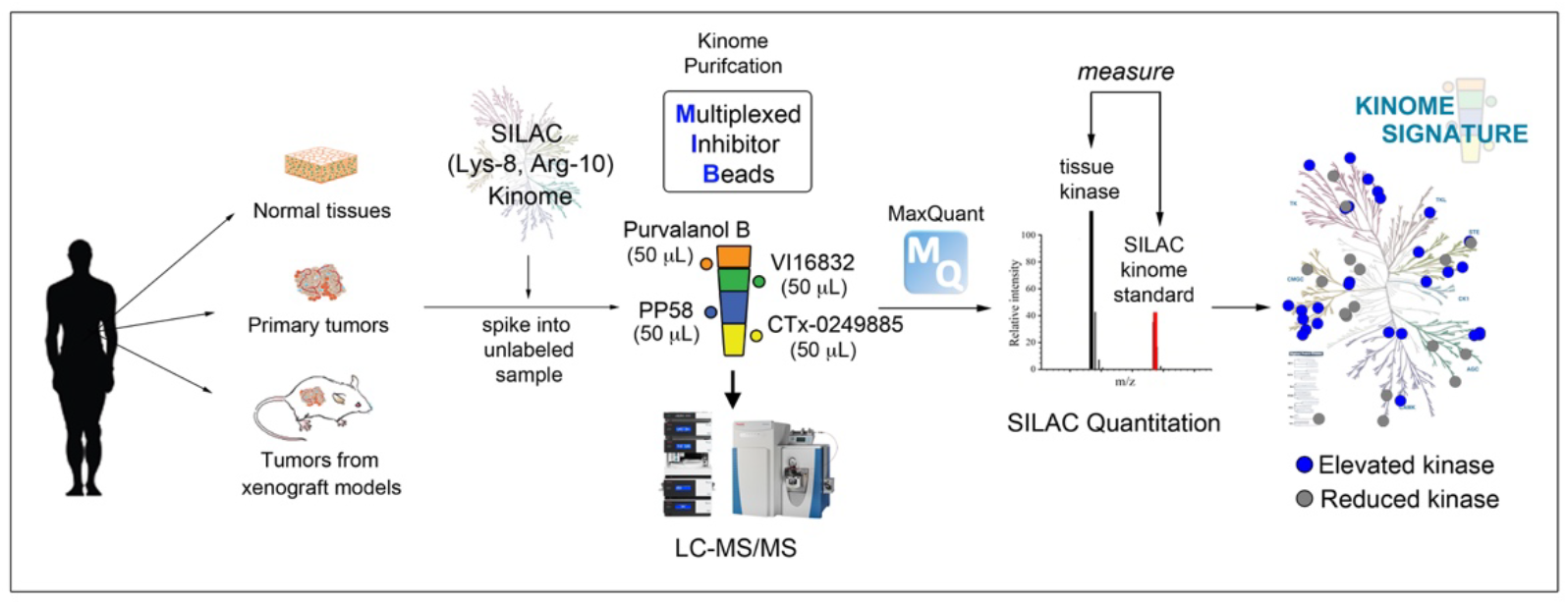
Workflow for measuring kinase abundance in tissues using Q-MIBs. An equal amount of our SILAC-based kinome standard (SKS) is added to any type of non-labeled sample (normal tissue, primary tumors or xenograft tumors) providing a robust proteomics strategy to quantify nearly 70% of the kinome across cohorts of samples, amongst treatments and across time.

To enrich endogenous protein kinases from tissue lysates, we utilized an equal mixture of 50 μL the pan-kinase inhibitor resins Purvalanol B, PP58, VI16832 and CTx-0249885, each inhibitor-resin purifying a distinct fraction of the kinome (**Fig. S1A, Data file S1**). CTx-0249885-resin exhibited the most diverse kinome recovery amongst inhibitor-beads. To quantify the kinome of tissue samples, we designed a super-SILAC kinome reference standard consisting of a cocktail of cancer cell lines that encompassed the majority of the human kinome. Eight cancer cell lines (UACC257, ACHN, OVCAR5, SF295, PC3, NCIH522, MOLT4, COLO205) were selected from the NCI-60 panel based on their diverse gene expression clustering (*26*), SILAC-labeled and kinase abundance measured by MIB-MS using the mixture of the unlabeled cell lines as a control reference (Fig. 2A). Hierarchical clustering and principal component analysis (PCA) of MIB-MS profiles revealed UACC257, MOLT4, COLO205, and ACHN exhibited the most distinct MIB-signatures, while OVCAR5, SF295, PC3, and NCIH522 shared more similar kinome profiles (Fig. 2B-C, and **S1B-C**, **Data file S1**). Following testing of various combinations, we identified the mixture of UACC257, MOLT4, COLO205, ACHN and PC3 cell lines provided the most diverse MIB-enriched kinome profile and was therefore selected as our SKS to be used repeatedly to quantify kinase levels across samples (Fig. 2D-E, **Data file S1**).

**Fig. 2.**
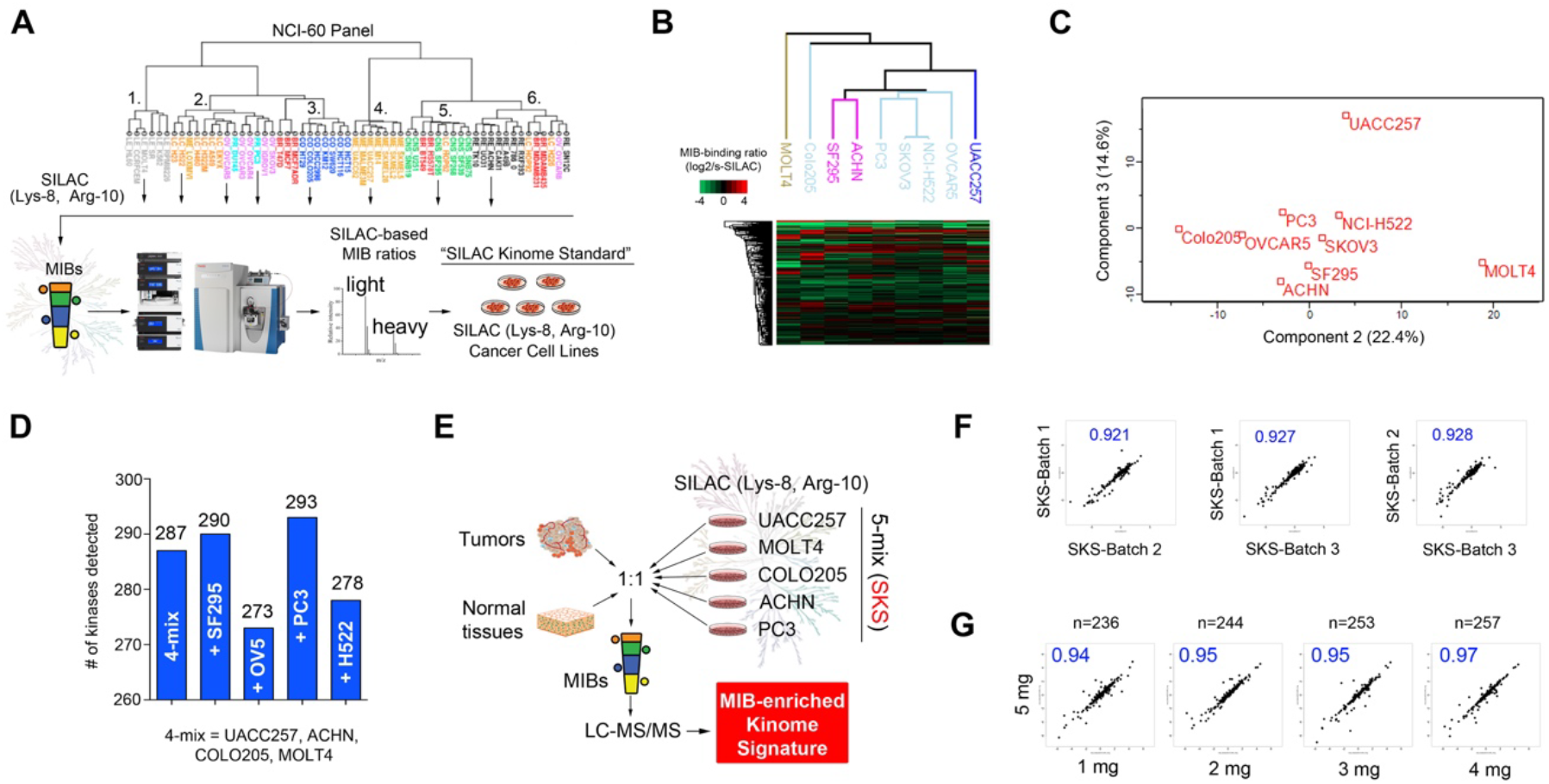
Designing a super-SILAC kinome standard for quantitating kinase levels in tissues using MIB-MS. A, 8 cancer cell lines were selected from the NCI-60 cell line panel to explore for the s-SILAC kinome standard based on published gene expression patterns (*26*). B, Hierarchical clustering of SILAC-determined kinase log2 ratios of the selected cancer cell lines. Each SILAC-labeled cell line was mixed 1:1 with the pooled non-labeled samples and analyzed by MIB-MS. C, Principal component analysis of SILAC-determined kinase log2 ratios of selected cancer cell lines. D, Number of kinases detected by MIB-MS profiling comparing various cell line mixtures. A mixture of the most diverse cell lines (UACC257, ACHN, COLO205 and MOLT4) was mixed 1:1 with SF295, OVCAR5, PC3 or NCI-H522 and kinome analyzed by MIB-MS. E, Selection of cancer cell lines that provided the most diverse MIB-enriched kinome to be used as the s-SILAC kinome standard for measuring kinases in tissues. F, Pearson correlation analysis of 3-distinct SKS batch preparations. Non-labeled OVCAR5 cells were mixed 1:1 with 3 distinct SKS batch preparations and kinome(s) measured by MIB-MS. G, Pearson correlation analysis of protein concentration inputs of samples. Various protein concentration of non-labeled OVCAR5 cells were mixed 1:1 with SKS and kinome(s) measured by MIB-MS.

The SKS can be propagated indefinitely, demonstrating high correlations in kinome quantitation across 3 distinct SKS batches prepared months apart, allowing quantitative comparison of kinome signatures from distinct kinome profiling projects (Fig. 2F, **Data file S1**). Next, we explored the optimal protein concentration input of the unlabeled sample to achieve maximal SILAC quantitation of the MIB-enriched kinome. The number of kinases measured by s-SILAC was overall similar amongst the various concentration of protein inputs ranging from 236 (1 mg) to 257 (5 mg), however, the number of kinases quantified by at least 3 unique peptides was lower using 1 mg input (n=173) compared to 5 mg protein input (n=201), favoring the use of 5 mg protein inputs if sample amounts are not limiting (**Fig. S1D, Data file S1)**. Various protein input concentrations produce highly correlated Q-MIB signatures, and although reduced numbers of kinases are quantitated using 1 mg relative to 5 mg inputs, those quantitated in both samples, exhibited high correlation (Fig. 2G). Thus, using the SKS permits the measurement and comparison of low yielding patient specimens such as biopsies with larger tumor sections from surgeries. Reproducibility of the MIB-MS paired with SKS assay to measure kinase abundance was confirmed using a model-based approach (*27*) for assessing technical reproducibility and outlier detection (**Fig. S1E**).

### Characterizing the Kinome Landscape Measured by Q-MIBs

Next, we explored the robustness of MIB-MS paired with SKS to measure kinase abundance in complex tissue samples. Measurement of kinases from tumors using Q-MIBs is highly reproducible across technical MIB replicates with R correlations values ≥0.9 comparing the same tumor kinome profiled on 3 separate days (Fig. 3A, **Data file S2**), as well as biological replicates of cell line xenograft tumors with R correlation values ~0.88 (Fig. 3B, **Data file S2**).

**Fig. 3.**
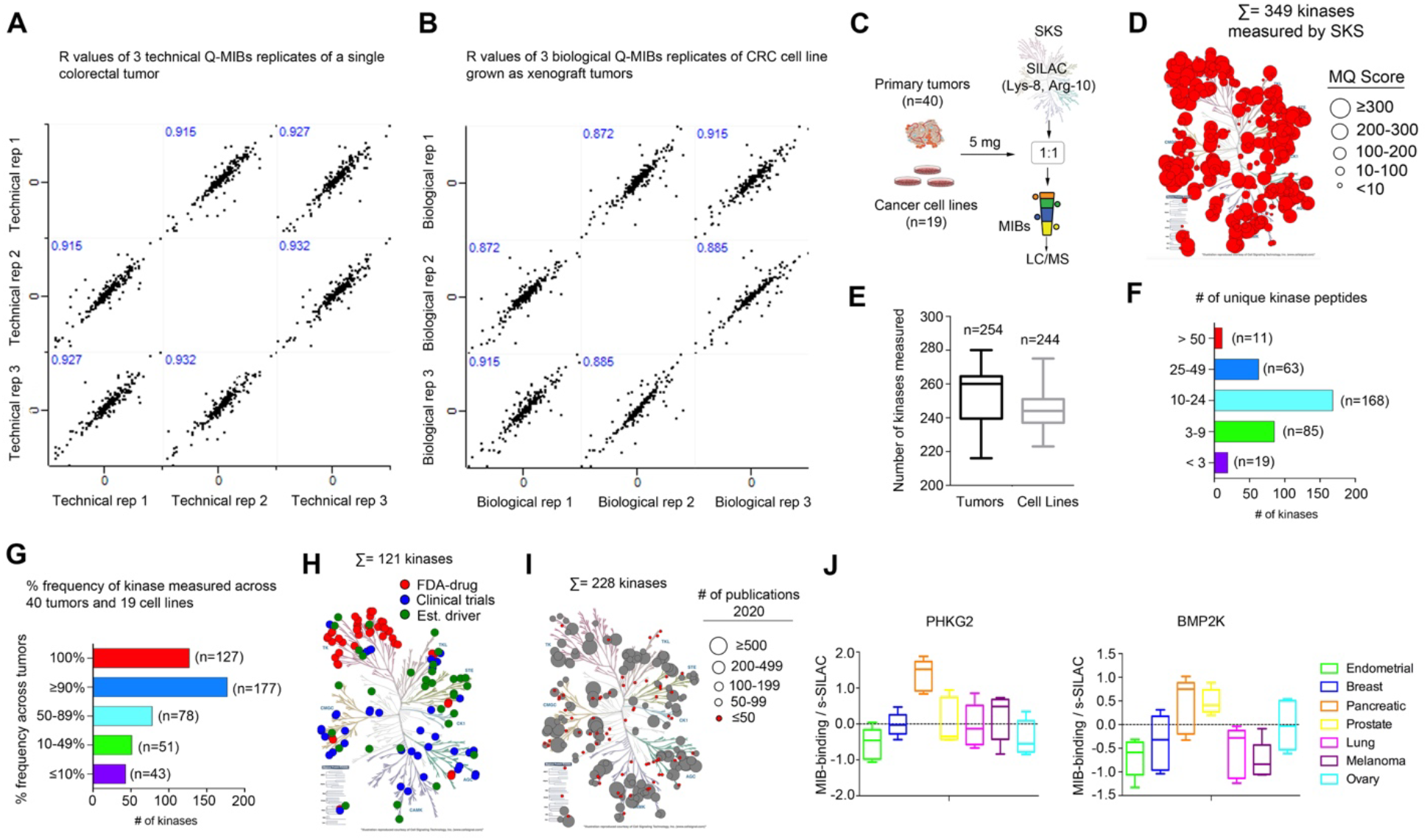
Characterization of the kinome measured by Q-MIBs across tissues. A, Pearson correlation analysis of MIB-MS profiles from 3-technical replicates of a single primary colorectal tumor. B, Pearson correlation analysis of 3-biological replicates of a tumor isolated from 3 distinct xenografts. The SW48 cell line was grown as a xenograft on 3-distinct mice, harvested and kinome(s) analyzed by MIB-MS. C, Study design for defining kinome measured by Q-MIBs. Lysates from snap frozen primary tumors from 8 distinct cancer types (prostate, breast, ovarian, endometrial, sarcoma, lung, melanoma, and pancreatic) and a collection of ovarian and endometrial cancer cell lines were mixed 1:1 with SKS and kinome(s) measured by MIB-MS. D, Kinome tree depicts fraction of kinome measured by Q-MIBs across all tumors and cells. Size of circle on tree represents MaxQuant score for MS/MS identification. Kinome tree was generated using KinMap. E, Box plot depicts the median number of kinases measured per run by Q-MIBs in tumors and cell lines. F, Bar plot shows the number of unique kinase peptides used to identify kinases across all tumor and cell lines. G, Bar plot depicts the percent frequency of kinases measured by MIB-MS across the 59 samples. H, Kinome tree depicts kinases measured by Q-MIBs with small molecules FDA-approved or drugs in clinical trial targeting the given kinases, as well as kinases with established driver functions (*5*). I, Kinome tree depicts fraction of the kinome measured by Q-MIBs not currently targeted for cancer therapy. Size of circles represent the number of publications for any given kinase. Pubmed central was search using keywords kinase names including aliases (2/1/20). J. Box plots show kinase abundance determined by Q-MIBs of understudied kinases PHKG2 and BMP2K amongst the various types of tumors. Five distinct tumors were kinome profiled from each tumor-type (n=5).

To define the fraction of the kinome measured by Q-MIBs, we kinome profiled a cohort of tumors (n=40) from 8 distinct cancer types, as well as a variety of cancer cell lines (n=19) using MIB-MS paired with the SKS (Fig. 3C, **Data file S2**). In total, 349 kinases were SILAC-quantitated with ~60% of the kinases exhibiting MQ scores >100 (Fig. 3D). The average number of kinases measured per sample was 254 (tumors) and 244 (cell lines) with the majority of kinases sequenced identified by 10-24 unique peptides (Fig. 3E-F, **Data file S2**). Frequency analysis of individual kinases measured across the 59 Q-MIBs samples showed 127 kinases were measured in every run, while 177 were measured at 90% frequency and ~73% were measured in ≥50% of the Q-MIBs runs (Fig. 3G, **Data file S2**). A detailed breakdown of individual kinase frequency of measurement by Q-MIBs across the 59 samples can be found in Data file S2.

Characterization of the kinome measured by Q-MIBs revealed significant coverage of kinases with FDA approved drugs (n=43), those currently being investigated in clinical trials (n=45) with several having established kinase driver function (n=33) (*5*) (Fig. 3H, **Data file S2**). It should be noted that most of the kinases quantified by Q-MIBs are largely untargeted for cancer therapies, with many (n=73) having sparse (<50) number of publications, representing the understudied or “dark” cancer kinome (Fig. 3I, **Data file S2**). For example, differential MIB-binding of understudied kinases PHKG2 and BMP2K was observed amongst tumor types, with pancreatic cancer exhibiting elevated levels of PHKG2, while BMP2K protein levels were the greatest in pancreatic and prostate tumors (Fig. 3J).

### Mapping the Kinome of Primary Endometrial Carcinoma Tumors Using Q-MIBs

Endometrial cancer (EC) is the most common gynecologic malignancy in the United States with 60,050 new cases and 10,470 deaths expected in 2020 (*28*). Most patients are diagnosed with uterine-confined disease (i.e. stage 1) (*28*) and thus, will have an overall favorable prognosis with 5 year disease-free survival greater than 80% (*29*). However, there has been a steady increase in the mortality rate for endometrial cancer which has been attributed to higher proportions of patients with advanced stage, higher grade and serous histology (*30*). Uterine Serous Carcinoma (USC) is one of the most common and lethal forms of endometrial carcinoma, and current treatments have only modestly impacted survival (*31*). To identify novel kinase therapeutic avenues in EC, we performed Q-MIBs profiling on 17 primary EC tumors (n=14 endometrioid, n=3 serous) and 14 normal endometrial (NE) tissues (Fig. 4A, **Fig. S2A and Data file S3**).

**Fig. 4.**
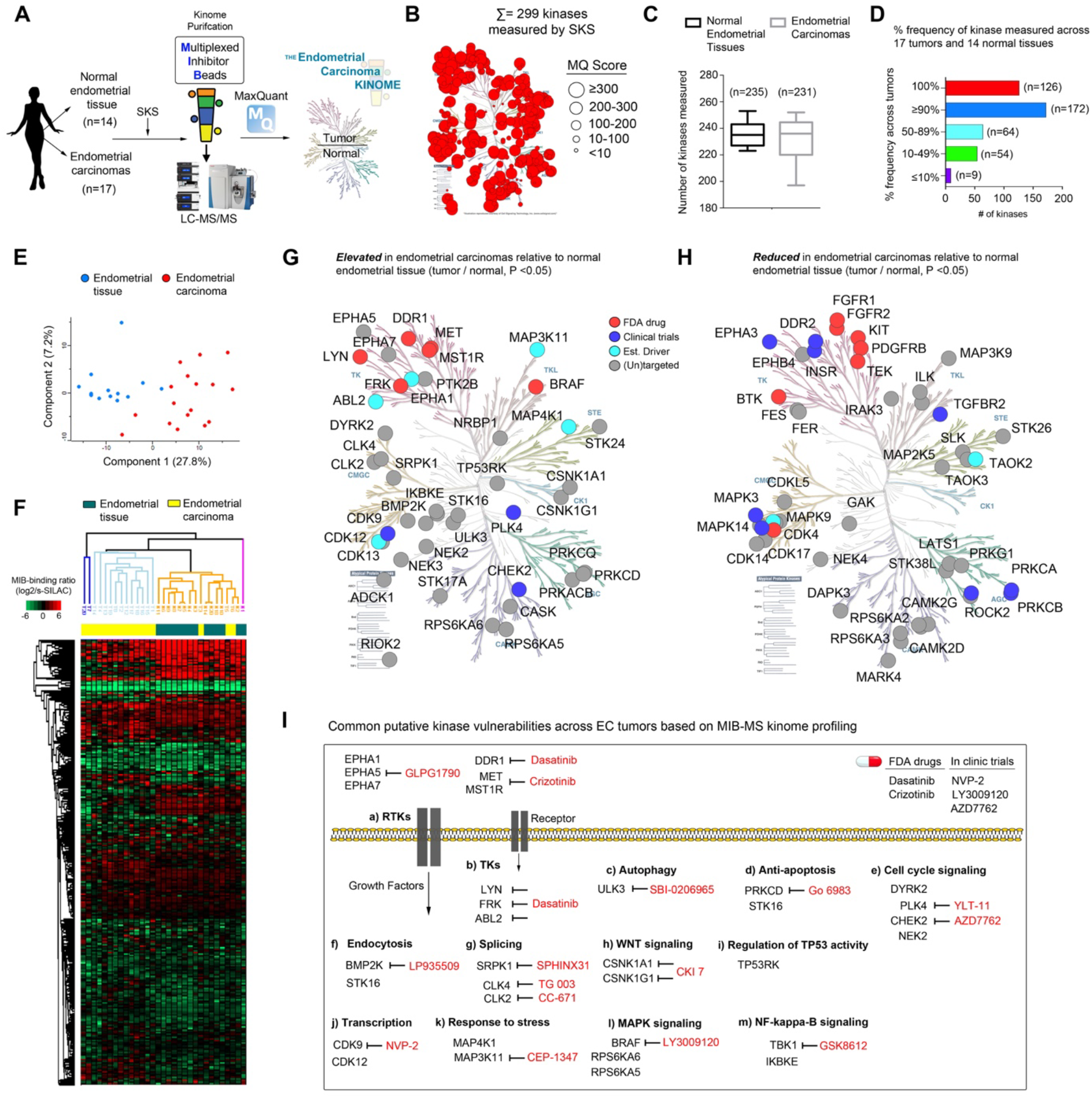
Profiling the kinome of primary endometrial carcinomas using Q-MIBs. A, Workflow for measuring kinase abundance in endometrial tumors and normal tissues using Q-MIBs. B, Kinome tree depicts fraction of kinome measured by Q-MIBs across all tumors and cells. Size of circle on tree represents MaxQuant score for MS/MS identification. C, Box plot depicts the median number of kinases measured by Q-MIBs in EC tumors and NE tissues. D, Bar plot depicts the percent frequency of kinases measured across all tumor and cell line Q-MIBs analysis. E, Principle component analysis of s-SILAC-determined kinase log2 ratios of EC tumors and NE tissues, as determined by MIB-MS. F, Hierarchical clustering of s-SILAC-determined kinase log2 ratios of EC tumors and NE tissues, as determined by MIB-MS. G-H, Kinome trees depict kinases statistical elevated (G) or reduced (H) in EC tumors relative to NE tissues. Statistical differences in kinase log2 s-SILAC ratios comparing EC tumors (n=17) vs NE tissues (n=14) was determined by paired *t*-test P < 0.05. I, Kinases and their associated pathways upregulated in EC tumors relative NE tissues. Available small molecule kinase inhibitors are denoted in red.

In total, we measured MIB-binding values for 299 kinases across EC tumors and NE tissues, with an average of 235 and 231 kinases measured per MIB-MS samples in NE tissues and EC tumors, respectively (Fig. 4B-C). Frequency analysis of kinase measurement across MIB-MS runs showed 126 kinases were measured in every run, while 236 kinases were quantitated in ≥50% of MIB-MS runs. (Fig. 4D, **Data file S3**). Principal component analysis or unsupervised hierarchical clustering of EC and NE tissue kinome profiles showed EC tumors clustered together and were overall distinct from NE tissues (Fig. 4E-F). Volcano plot analysis of MIB-MS signatures amongst EC tumors and NE tissues revealed many kinases were differentially expressed (**Fig. S2B**). Kinases commonly elevated (Fig. 4G) or reduced (Fig. 4H) in tumors relative to NE tissues are depicted on kinome trees.

Many kinases overexpressed in EC tumors have inhibitors FDA-approved inhibitors or drugs currently being evaluated in clinical trials, representing potentially actionable kinases for treatment (Fig. 4I). The pan-Tyrosine kinase (TK) inhibitor Dasatinib, represents a plausible drug therapy for EC tumors based on increased MIB-binding of Dasatinib-targets LYN, DDR1 and/or SRC. Indeed, Dasatinib is currently being evaluated in EC highlighting the utility of Q-MIBs to predict kinase drug targets (*32*). Elevated levels of the RTK, c-MET (MET), were also observed in EC tumors relative to NE tissues by Q-MIBs profiling, consistent with previous reports demonstrating high expression of c-MET and its ligand hepatocyte growth factor (HGF) in endometrial carcinomas (*33*). Several FDA-approved kinase inhibitors that target MET are available, such as Crizotinib and several groups have shown blockade of MET signaling impairs endometrial cancer cell growth and invasion (*34*). Elevated MIB-binding of CDK9 was detected in EC tumors supporting the use of CDK9 inhibitors for the treatment of EC. CDK9 has been shown to be an essential component of the P-TEF complex regulating transcriptional elongation, where inhibition of CDK9 results in blockade of transcription of oncogenes such as MYC (*35*). Of particular interest, several of the kinases overexpressed in EC tumors represented understudied kinases with no previous association with EC. Moreover, high quality kinase inhibitors were available for a number of these kinases, including CLK2, ULK3, BMP2K and SRPK1, making these attractive targets for subsequent loss-of-function studies in EC cell lines.

### SRPK1 Is Overexpressed in Endometrial Tumors and Represents a Prognostic Indicator for Poor Survival

Serine/Arginine-Rich Splicing Factor kinase, (SRPK1) overexpression has been correlated with poor survival of several cancers including colorectal, lung and breast cancers (*36*). Overexpression of SRPK1 has been observed in highly aggressive breast cancers, such as triple negative breast cancers (TNBCs) and may be linked to development of resistance to therapy, highlighting SRPK1 as an attractive therapeutic target (*37*). SRPK1 has been shown to be essential for mRNA alternative splicing in cells and blockade of SRPK1 function alters several oncogenic processes including angiogenesis, migration/invasion, proliferation, cancer stem cell (CSC) phenotype, and sensitivity to chemotherapy (*38*).

In the present study, Q-MIBs kinome profiling identified SRPK1 to be elevated in EC tumors relative to normal endometrial tissues (Fig. 5A). Immunoblot analysis comparing SRPK1 protein levels amongst EC tumors and paired matched normal tissue confirmed Q-MIBs findings, where the majority of EC tumors overexpressed SRPK1 (Fig. 5B). Notably, both EC tumor subtypes, endometrioid or uterine serous carcinoma (USC) displayed elevated levels of SRPK1 compared to NE tissues by immunoblot analysis. Elevated protein levels of SRPK1 in EC was confirmed by immunohistochemical (IHC) analysis of tissue microarrays (TMAs) containing 57 EC patient tumors, (39 USC and 18 endometrioid) and 12 normal endometrial tissues (Fig. 5C-D and **Data file S4**).

**Fig. 5.**
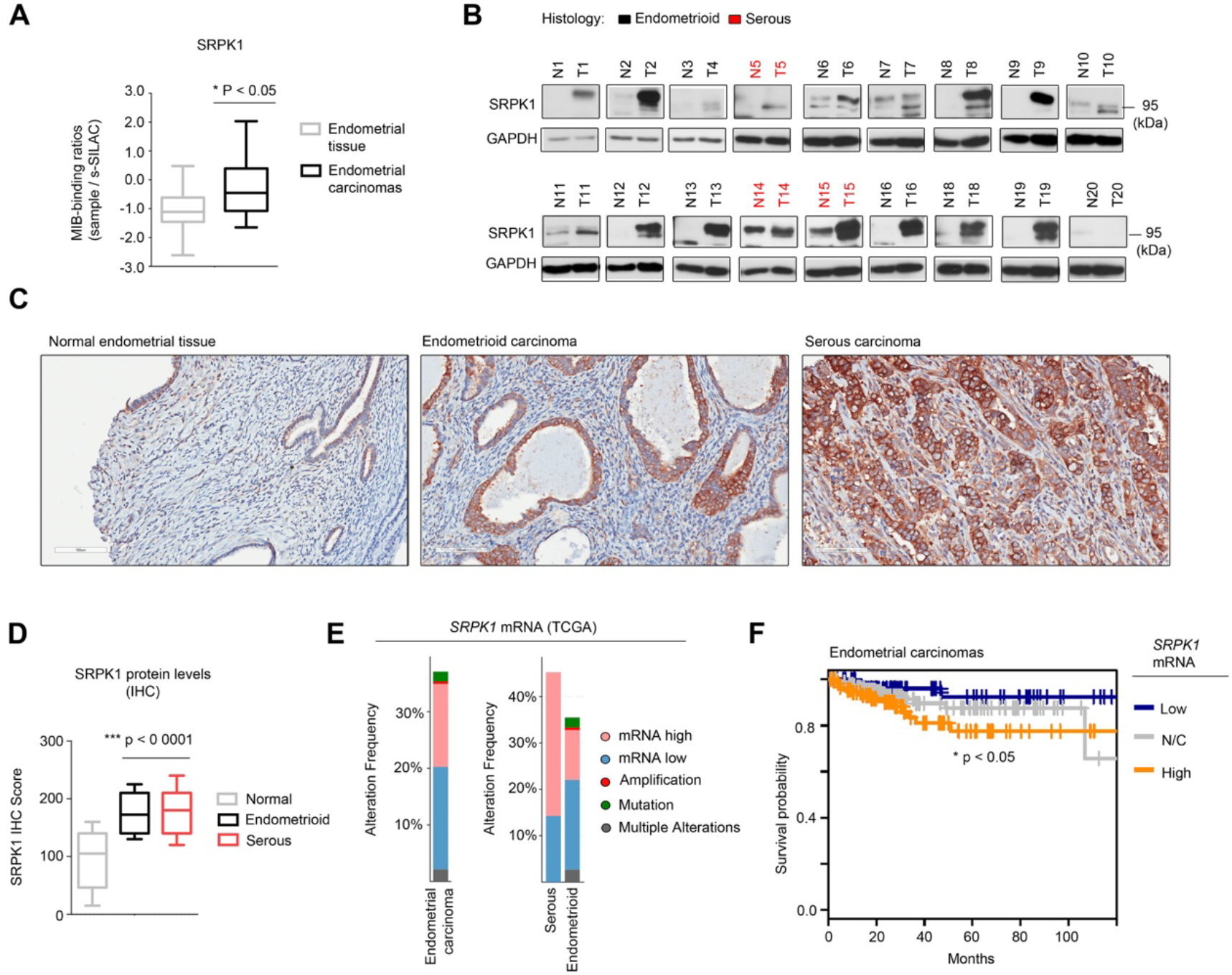
SRPK1 is overexpressed in EC tumors and associated with poor survival. A, Box plot depicts elevated SRPK1 levels in EC tumors relative to NE tissues as determined by Q-MIBs. Statistical differences in SRPK1 log2 s-SILAC ratios comparing EC tumors (n=17) vs NE tissues (n=14) was determined by paired *t*-test P < 0.05. B, Immunoblot analysis of SRPK1 protein abundance comparing EC tumors (n=20) and matched normal adjacent tissues used in the Q-MIBs analysis. C, IHC analysis of SRPK1 in EC TMAs. Immunoreactivity of SRPK1 protein was evaluated by a pathologist and IHC H score was given based on the intensity and percentage of protein stain on tumor cells. SRPK1 IHC images captured at 100 um. D, Box plots shows elevated SRPK1 protein abundance in endometrioid and serous subtypes of EC tumors relative to NE tissues as determined by IHC. Data are from duplicate analysis of TMAs consisting of 57 EC tumors (39 USC and 18 endometrioid) and 12 NE tissues. Statistical differences in SRPK1 IHC scores comparing EC tumors vs NE tissues was determined by paired *t*-test P < 0.05. E, Analysis of *SRPK1* copy number analysis (CNA), mRNA and mutation changes in EC tumors from TCGA studies (*17*). mRNA levels determined by U133 microarray and change in *SRPK1* abundance amongst tumors determined at z-score > 1 or < −1. F, Survival plot of HGSOC patients with *SRPK1* expression segregated into lower (low-blue), middle (no change – grey) and upper quartiles (Up; orange) show statistical difference in overall survival. Survival data was obtained from TCGA data (*17*).

To characterize SRPK1 expression across larger numbers of patient EC tumors, we queried publicly available EC genomics studies (*17*). Genomic alteration of *SRPK1* (37%, 86 of 232 EC tumors) was observed in EC tumors, with frequent altered gene expression (upregulation in 15%, 34 of 232 and downregulation in 18%, 42 of 232). Infrequent gene amplification (0.43%, 1 of 232) and mutations in *SRPK1* were observed (1.72%, 4 of 232, with no homozygous deletions detected (Fig. 5E). Notably, *SRPK1* mRNA levels were upregulated in 31% of USC tumors (13 out of 42) and 11% of endometrioid tumors (20 out of 186). Levels of *SRPK1* mRNA expression were associated with survival in EC, where patients harboring EC tumors with high levels of *SRPK1* exhibited reduced overall survival (P <0.05) (Fig. 5F).

### SRPK1 Inhibition Promotes Alternative Splicing Impairing Survival Signaling in USC Cells

SRPK1 has established roles in the regulation of alternative splicing in cancer (*39*). To explore the role of SRPK1 in control of mRNA splicing in EC cells, we treated an established USC cell line SPEC-2, with the highly selective SRPK1 inhibitor, SPHINX31 (*40*) and assessed the impact on alternative splicing. Previous reports showed inhibition of SRPK1 in serum-starved cells induced significant alternative splicing of VEGF-A (*40, 41*). Therefore, we assessed the impact of SPHINX31-treatment on mRNA splicing in serum-starved SPEC-2 cells. Phosphorylation of SRSF (SR Splicing Factors) by SRPK1 has been shown to regulate splicing, RNA export, and other processes of RNA metabolism in cells. Treatment of SPEC-2 cells with SPHINX31 reduced phosphorylation of all major forms of SRPK1-substrate SRSF (SR Splicing Factors), demonstrating SRPK1 inhibition blocks phosphorylation of splicing factors (Fig. 6A). To gain insight into the role of SRPK1 in alternative splicing in EC, we performed RNA-seq analysis on SPEC-2 cells treated with SPHINX31 for 24 h and assessed changes in pre-mRNA splicing. RNA-seq analysis revealed SRPK1 inhibition resulted in substantial mis-splicing of gene targets, including 2759 SKIP events, 518 mutually exclusive exons (MXE), 225 5’ splice site (5’SS), 364 3’ splice site (3’SS) and 640 retained intron (RI) (Fig. 6B, **Data file S5**). Pathway analysis of genes exhibiting mis-splicing following SRPK1 inhibition were enriched in cell-cell adhesion, protein transport, cell cycle, protein phosphorylation and mRNA splicing signaling (Fig. 6C). Similarly, BIOCARTA pathway analysis showed an enrichment of alternative spliced targets in RHO pathway, ECM pathway and VEGF pathway signaling (Fig. 6D). Consistent with these findings, inhibition of SRPK1 has been previously shown to block splicing of VEGF 165a to VEGF 165b impacting Vegf signaling and angiogenesis (*38*).

**Fig. 6.**
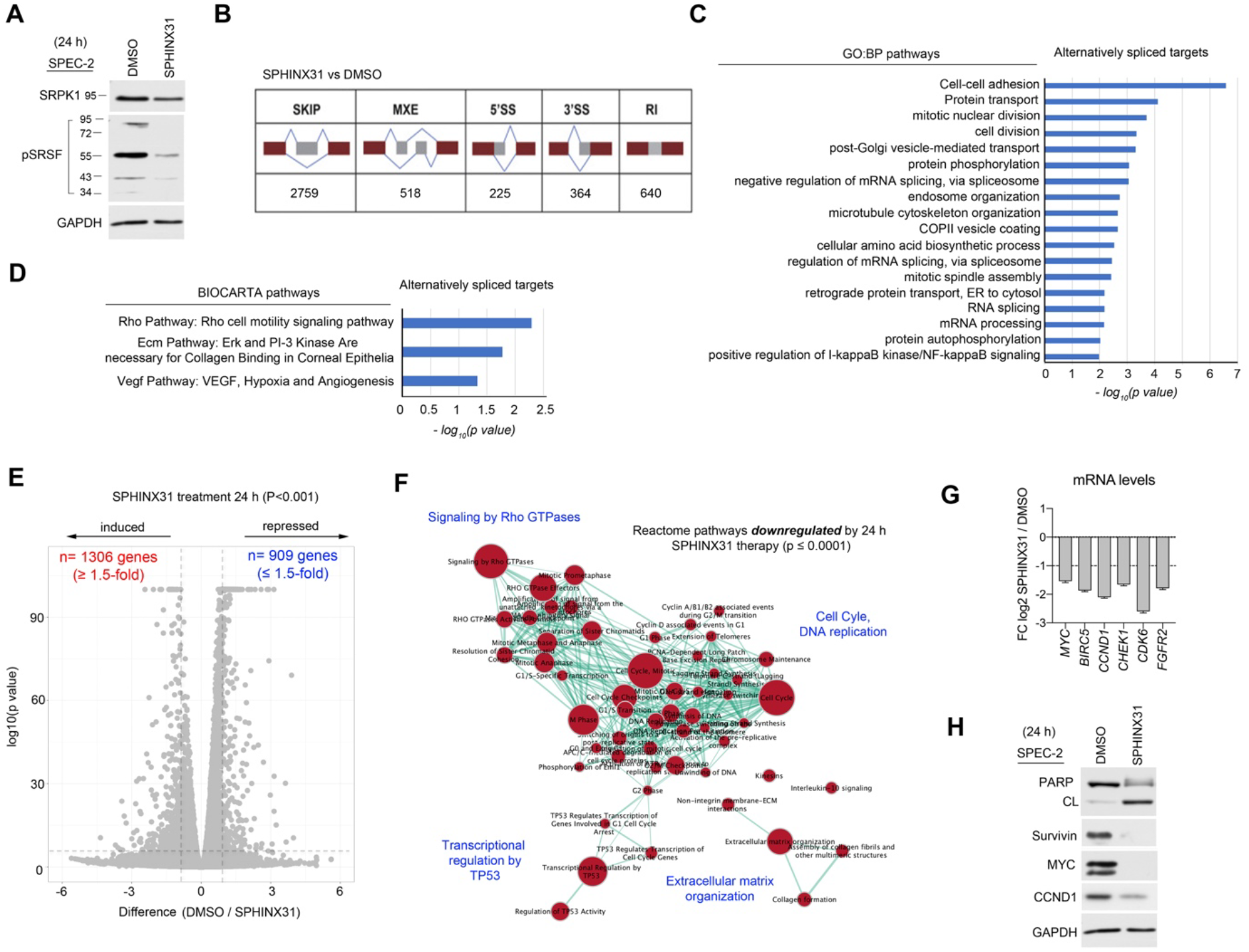
Inhibition of SRPK1 induces alternative splicing in USC cells causing apoptosis. A, Immunoblot of serum-starved SPEC-2 cells treated with SPHINX31 (5 μM) for 24 h. B, Illustration showing the type of alternative splicing events identified using RNA-seq between SPHINX3-treatment and controls with number of events at exons shown below. C, GO Biological Process pathways analysis of transcripts affected by SPHINX31-treatment from RNA-seq analysis was performed using Database for Annotation, Visualization and Integrated Discovery (DAVID) v6.8. Significant GO Biological Process terms (P <0.05) are depicted as a bar graph with the p values represented as -log10 on the x axis. D, BIOCARTA pathways analysis of transcripts affected by SPHINX31-treatment from RNA-seq analysis. Significant BIOCARTA terms (P <0.05) are depicted as a bar graph with the p values represented as -log10 on the x axis. E, Volcano plot depicts log_2_fold changes in gene expression as a ratio of DMSO / SPHINX31. SPEC-2 cells were treated with DMSO or SPHINX31 (5 μM) for 24 h and changes in gene expression assessed by RNA-seq. Statistical differences in gene expression comparing SPHINX31-treated and control treated cells was determined by paired T-Test Benjamini-Hochberg adjusted p values at FDR of <0.05. F, Enrichment map of pathways repressed by SPHINX31 treatment (5 μM) for 24 h based on RNA-seq analysis. Pathway enrichment maps were generated in Cytoscape using the Enrichment Map app v2.2.1. G, Bar plot depicts established oncogenes whose mRNA levels were reduced by 24 h SPHINX31 (5 μM) treatment in SPEC-2 cells. Changes in mRNA levels were determined by RNA-seq. H, Immunoblot confirms reduced MYC, Survivin and Cyclin D1 protein levels in response to 24 h SPHINX31 (5 μM) treatment.

Analysis of mRNA expression changes following SPHINX31-treatment of SPEC-2 cells revealed SRPK1 inhibition induced significant changes to gene expression, including 1306 gene upregulated and 909 genes downregulated (Fig. 6E, **Data file S5**). Reactome pathway analysis of genes downregulated by SPHINX31-treatment were enriched in Rho GTPase, cell cycle, DNA replication, transcriptional regulation by TP53 and extracellular matrix organization signaling (Fig. 6F and **S3A**). Genes downregulated following SRPK1 inhibition were also enriched in PI3K-AKT and FoxO signaling, consistent with previous reports demonstrating SRPK1 inhibition impaired AKT signaling through disruption with PHLPP interaction (*42*) (**Fig. S3B**). Moreover, several genes involved in negative regulation of apoptosis and cell cycle progression were downregulated in response to SRPK1 blockage, including *BIRC5* (Survivin), *MYC*, *MYCN* and *CCND1*, all established oncogenic drivers (Fig. 6G). Knockdown of Survivin has been shown to induce apoptosis and mitotic catastrophe in several cancer cell lines (*43*), while MYC depletion induces cell cycle arrest (*44*). Immunoblot analysis of SPHINX31-treated cells confirmed several mRNA expression changes including reduced protein levels of established oncogenes, MYC and Survivin (Fig. 6H). Furthermore, an increase in cleaved PARP was observed by immunoblot, demonstrating SRPK1 inhibition induces apoptosis in SPEC-2 cells.

Together, SRPK1 inhibition inhibited cell viability and induced apoptosis distinctly in serum-starved USC cells. Moreover, SPHINX31-treatment induced significant alternative splicing of genes involved in cell motility and cell cycle resulting in the downregulation of several genes essential for growth and survival, including oncogenes MYC and Survivin. Finally, SPHINX31-treatment induced apoptosis in USC cells nominating SRPK1 as a plausible drug target for the treatment of EC.

## DISCUSSION

The protein kinome represents one of the most promising and actionable classes of drug targets for the treatment of cancer; yet, the majority of the kinome remains untargeted for drug therapy, with many kinases having no established oncogenic function (*4, 6*). The vast majority of kinase publications focus on a small group of well-understood kinases, yet synthetic lethal screens repeatedly identify various untargeted kinases as playing a vital role in cancer cell proliferation and survival (*45*). Our inability to routinely probe these enzymes has hindered previous attempts to understand how they are regulated and function in cancer. Importantly, emergence of unbiased proteomics strategies such as Kinobeads and MIB-MS has provided methods to capture and quantify these elusive untargeted kinases in normal and in cancer cells (*12, 46*). Notably, MIB-MS has been extensively utilized in the study of cancer cell line models, however, its application in tumor models has not been systematically characterized. Here, we developed a SILAC-based MIB-MS platform (Q-MIBs) specifically designed to profile the kinome of human tissues, providing a quantitative assay for measuring kinase abundance of nearly 70% of the kinome. Application of Q-MIBs in endometrial cancer identified a network of kinases overexpressed in EC tumors, including SRPK1, a kinase involved in mRNA splicing, previously unexplored in EC. Subsequent proteogenomics and loss-of-function studies demonstrated overexpression of SRPK1 was correlated with poor survival in EC and inhibition of SRPK1 in a USC cell line induced significant alternative splicing promoting apoptosis.

Several methods for quantitating kinases enriched by MIBs or kinobeads have been explored including isobaric tagging (i.e., iTRAQ or TMT), label-free quantitation (LFQ) approaches, and labelled methods such as stable isotope labeling with amino acids in cell culture (SILAC), each with strengths and weaknesses. Several groups have reported ratio compression arising from contamination during precursor ion selection in MS2 quantitation using isobaric tagging methods, leading to compromised accuracy, compression and underestimation of the ratios (*47, 48*). A potential drawback to label-free quantitation is that experimental conditions such as variations in kinase enrichment from MIB-columns amongst samples, LC/MS run conditions such as temperature, and column conditions may present differences between samples (*49*). Using labelling techniques such as SILAC or super-SILAC has the benefit of measuring samples within the same MS run (*20, 24*). Here, we developed a super-SILAC kinome standard to spike into tissue samples to control for variations in kinase MIB-binding and/or LC-MS/MS retention time reproducibility. However, as SILAC approaches measure protein abundance by MS1 quantitation, they can result in lower overall protein coverage (*50*). Thus, a potential limitation to measuring kinase levels in heterogeneous tissues using the Q-MIBs approach is that we can only quantitate with high confidence those kinases that have a matching SILAC-paired kinase in SKS, if the kinase is present at low levels or not expressed in the SKS cell lines, we could potentially miss kinase measurements in tissue samples. Recently, it has been shown that using a SILAC-labelled control sample mixed with each unlabeled sample prior to MS analysis, (termed-QuantFusion) can help control for experimental and/or instrumental variations amongst samples observed with LFQ approaches (*51*). Importantly, as tissues samples profiled by Q-MIBs are unlabeled, we could re-analyze the MS data using MaxQuant software quantitating only the non-labeled kinase peptides by MaxLFQ (*52*). Combining s-SILAC and LFQ could provide a complementary quantitation strategy to confirm MIB-MS determined kinase abundance measurements, as well as to measure the levels of kinases not preset in the SKS using LFQ, improving coverage of the kinome measured in tissues.

Uterine Serous Carcinoma (USC) is one of the most common and lethal forms of endometrial carcinoma, and current treatments have only modestly impacted survival (*31*). These tumors are characterized by alterations in p53, p16, E-cadherin, ERBB2, PIK3CA, FBXW7, and PPP2R1A (*53, 54*), and consequently, exhibit genome instability and aberrant signaling. Signaling abnormalities in USCs, particularly those involving over-activated protein kinases, represent potential therapeutic avenues. Here, we showed inhibition of protein kinase SRPK1 significantly altered mRNA splicing resulting apoptosis in an established USC cell line, nominating SRPK1 as a plausible therapeutic target for USC. Inhibition of SRPK1 in USC cells altered splicing of genes involved in a variety of oncogenic processes including cell migration, cell cycle and apoptosis. Mis-spliced gene targets following SPHINX31-treatment were enriched in VEGF pathways and angiogenesis, consistent with recent studies showing SRPK1 function has been shown to block splicing of VEGF 165a to VEGF 165b reducing angiogenesis in cells and tumor models (*38*). Importantly, Bevacizumab, an antibody against VEGF-A has shown moderate clinical responses in EC, suggesting SRPK1 inhibition could represent a promising anti-angiogenic therapy for EC (*16*). However, further studies in tumor models will be required to determine the impact of SRPK1 inhibition on angiogenesis and EC tumor growth. Recently, SPRK1 was shown to regulate the splicing of BRD4 in AML cells, where SPHINX31-treatment promotes the splicing of the long form of BRD4 (*55*). BRD4 has been shown to act as enhancer of MYC transcription in several cancer models, and inhibition of BRD4 blocks MYC transcription resulting in tumor apoptosis. Here, we observed a reduction in MYC and MYCN at the RNA and protein level in response to SRPK1 inhibition, suggesting BRD4 could be impacted by SPHINX31 treatment, however, a detailed analysis of BRD4 splicing will be required. Notably, BET inhibitors have recently been shown to be effective at inhibiting growth of MYC-overexpressed USC cells and tumors (*56*), suggesting SPRK1 blockade may represent a therapeutic option for MYC-driven USCs.

Recently, anti-PD-1/PD-L1 antibody checkpoint inhibitors have emerged as a promising immunotherapy for the treatment of endometrial cancer (*57*). Moreover, FDA granted accelerated approval to the combination of checkpoint inhibitors and the VEGFR inhibitor lenvatinib for the treatment of patients with advanced endometrial cancer with disease progression following prior systemic therapy (*58*). Several studies have shown that peptides derived from altered mRNA splicing in tumors could bind to MHC class I molecules representing tumor-specific mRNA splicing derived neoantigens (*59*). Notably, inhibition of SRPK1 induced > 1600 mis-spliced events in USC cells, suggesting treatment of USC with SRPK1 inhibitors could enhance immunoreactivity of USC tumors improving anti-PD1/PD-L1 antibody checkpoint inhibitors. Future studies characterizing SRPK1-inhibitor induced mRNA splicing and its impact on immunoreactivity in EC tumor models will be of particular interest.

Together, probing the endometrial carcinoma tumor kinome using a newly designed SILAC-based MIB-MS platform led to the discovery of SRPK1 as an essential kinase regulating mRNA splicing and survival of EC cells, highlighting SPRK1 as a plausible drug target for the treatment of EC.

## Acknowledgements

We thank Chunxiao Zhou (University of North Carolina) for providing the SPEC-2 cell line.

## Author contributions

J.S.D. wrote the manuscript. A.M.K. performed all western blots. N.S. performed western blots and tumor xenograft experiments. A.M.K. performed all cell culture experiments. K.J.J., V.K., and A.M.K. performed all MIB-MS experiments. V.K., and N.S. processed all tumors and tissues. S.P. performed RNA sequencing, splicing studies and bioinformatics analysis of TCGA datasets. K.Q.C. carried out IHC analysis of EC tumors. J.S.D., K.J.J., V.K., S.P., and A.M.K., contributed to experimental design.

## Data and Materials Availability

The mass spectrometry proteomics data have been deposited to the ProteomeXchange Consortium via the PRIDE partner repository with the dataset identifier PXD017785. The BioProject ID number for the mRNA sequencing data reported in this paper is PRJNA608466. All other data needed to evaluate the conclusion in the paper are present in the paper or the Supplementary Materials.

## SUPPLEMENTARY MATERIALS

Figure S1: Designing a s-SILAC kinase standard for measuring kinase abundance in tissues.

Figure S2: Kinome profiling of EC tumors and NE tissues using Q-MIBs.

Figure S3. Pathway analysis of genes downregulated by SRPK1 inhibition in USC cells.

Data file S1: Design and testing of s-SILAC kinome reference standard for MIB-MS profiling of tissues. Tables relate to Figure 2 and Figure S1 displaying raw unique peptides and s-SILAC ratios from MIB-MS profiling experiments.

Data file S2. Characterizing the kinome measured by Q-MIBs across a cohort of tissues and cell lines. Tables relate to Figure 3 displaying raw unique peptides and s-SILAC ratios from MIB-MS profiling experiments.

Data file S3. Mapping the kinome landscape of endometrial tumors and normal endometrial tissues using Q-MIBs.

Data file S4. Immunohistochemistry of SRPK1 in endometrial tumors and normal endometrial tissues using tissue microarrays (TMAs).

Data file S5. SRPK1 inhibition alters splicing and gene expression in SPEC-2 cells

## REFERENCES

1. G. Manning, D. B. Whyte, R. Martinez, T. Hunter, S. Sudarsanam, The Protein Kinase Complement of the Human Genome. Science 298, 1912–1934 (2002).

2. J. T. Metz et al., Navigating the kinome. Nat. Chem. Biol. 7, 200–202 (2011).

3. Z. A. Knight, H. Lin, K. M. Shokat, Targeting the cancer kinome through polypharmacology. Nat. Rev. Cancer 10, 130–137 (2010).

4. S. Klaeger et al., The target landscape of clinical kinase drugs. Science 358, (2017).

5. E. D. G. Fleuren, L. Zhang, J. Wu, R. J. Daly, The kinome ‘at large’ in cancer. Nat. Rev. Cancer 16, 83–98 (2016).

6. O. Fedorov, S. Muller, S. Knapp, The (un)targeted cancer kinome. Nat. Chem. Biol. 6, 166–169 (2010).

7. S. Knapp et al., A public-private partnership to unlock the untargeted kinome. Nat. Chem. Biol. 9, 3–6 (2013).

8. D. H. Drewry et al., Progress towards a public chemogenomic set for protein kinases and a call for contributions. PLoS One 12, e0181585 (2017).

9. C. Scholl et al., Synthetic Lethal Interaction between Oncogenic KRAS Dependency and STK33 Suppression in Human Cancer Cells. Cell 137, 821–834 (2009).

10. D. A. Barbie et al., Systematic RNA interference reveals that oncogenic KRAS-driven cancers require TBK1. Nature 462, 108–112 (2009).

11. M. Bantscheff et al., Quantitative chemical proteomics reveals mechanisms of action of clinical ABL kinase inhibitors. Nat. Biotechnol. 25, 1035–1044 (2007).

12. James S. Duncan et al., Dynamic Reprogramming of the Kinome in Response to Targeted MEK Inhibition in Triple-Negative Breast Cancer. Cell 149, 307–321 (2012).

13. C. E. Franks, K.-L. Hsu, Activity-Based Kinome Profiling Using Chemical Proteomics and ATP Acyl Phosphates. Curr. Protoc. Chem. Biol. 11, e72–e72 (2019).

14. L. M. Graves, J. S. Duncan, M. C. Whittle, G. L. Johnson, The dynamic nature of the kinome. Biochem. J. 450, 1–8 (2013).

15. H. Gao et al., High-throughput screening using patient-derived tumor xenografts to predict clinical trial drug response. Nat. Med. 21, 1318–1325 (2015).

16. L. M. Charo, S. C. Plaxe, Recent advances in endometrial cancer: a review of key clinical trials from 2015 to 2019. F1000Res 8, F1000 Faculty Rev-1849 (2019).

17. N. Cancer Genome Atlas Research et al., Integrated genomic characterization of endometrial carcinoma. Nature 497, 67–73 (2013).

18. C. Trapnell et al., Differential gene and transcript expression analysis of RNA-seq experiments with TopHat and Cufflinks. Nat. Protocols 7, 562–578 (2012).

19. S. Shen et al., MATS: a Bayesian framework for flexible detection of differential alternative splicing from RNA-Seq data. Nucleic Acids Res. 40, e61–e61 (2012).

20. B. B. Ong SE, Kratchmarova I, Kristensen DB, Steen H, Pandey A, Mann M., Stable isotope labeling by amino acids in cell culture, SILAC, as a simple and accurate approach to expression proteomics. Mol. Cell. Proteomics 1, 376–386 (2002).

21. A. M. Kurimchak et al., Resistance to BET Bromodomain Inhibitors Is Mediated by Kinome Reprogramming in Ovarian Cancer. Cell reports 16, 1273–1286 (2016).

22. Y. Zhang et al., Ribosomal Proteins Rpl22 and Rpl22l1 Control Morphogenesis by Regulating Pre-mRNA Splicing. Cell reports 18, 545–556 (2017).

23. S. Falcon, R. Gentleman, Using GOstats to test gene lists for GO term association. Bioinformatics 23, 257–258 (2006).

24. T. A. Neubert, P. Tempst, Super-SILAC for tumors and tissues. Nat Meth 7, 361–362 (2010).

25. J. Cox et al., A practical guide to the MaxQuant computational platform for SILAC-based quantitative proteomics. Nat. Protoc. 4, 698–705 (2009).

26. Amin M. Gholami et al., Global Proteome Analysis of the NCI-60 Cell Line Panel. Cell Reports 4, 609–620 (2013).

27. T. Anastassiadis, S. W. Deacon, K. Devarajan, H. Ma, J. R. Peterson, Comprehensive assay of kinase catalytic activity reveals features of kinase inhibitor selectivity. Nat Biotech 29, 1039–1045 (2011).

28. R. L. Siegel, K. D. Miller, A. Jemal, Cancer statistics, 2016. CA Cancer J Clin 66, 7–30 (2016).

29. H. Lajer et al., Survival after stage IA endometrial cancer; can follow-up be altered? A prospective nationwide Danish survey. Acta Obstet Gynecol Scand 91, 976–982 (2012).

30. S. M. Ueda et al., Trends in demographic and clinical characteristics in women diagnosed with corpus cancer and their potential impact on the increasing number of deaths. Am J Obstet Gynecol 198, 218 e211-216 (2008).

31. M. G. del Carmen, M. Birrer, J. O. Schorge, Uterine papillary serous cancer: a review of the literature. Gynecol. Oncol. 127, 651–661 (2012).

32. L. R. Duska et al., A window-of-opportunity clinical trial of dasatinib in women with newly diagnosed endometrial cancer. Cancer Chemother. Pharmacol. 83, 473–482 (2019).

33. M. Li, X. Xin, T. Wu, T. Hua, H. Wang, HGF and c-Met in pathogenesis of endometrial carcinoma. Front Biosci (Landmark Ed) 20, 635–643 (2015).

34. Y. Kogata et al., Foretinib (GSK1363089) induces p53-dependent apoptosis in endometrial cancer. Oncotarget 9, 22769–22784 (2018).

35. F. Morales, A. Giordano, Overview of CDK9 as a target in cancer research. Cell cycle (Georgetown, Tex.) 15, 519–527 (2016).

36. N. Bullock, S. Oltean, The many faces of SRPK1. The Journal of pathology 241, 437–440 (2017).

37. W. van Roosmalen et al., Tumor cell migration screen identifies SRPK1 as breast cancer metastasis determinant. The Journal of clinical investigation 125, 1648–1664 (2015).

38. N. Bullock et al., Serine-arginine protein kinase 1 (SRPK1), a determinant of angiogenesis, is upregulated in prostate cancer and correlates with disease stage and invasion. J. Clin. Pathol. 69, 171–175 (2016).

39. A. Czubaty, A. Piekiełko-Witkowska, Protein kinases that phosphorylate splicing factors: Roles in cancer development, progression and possible therapeutic options. The international journal of biochemistry & cell biology 91, 102–115 (2017).

40. J. Batson et al., Development of Potent, Selective SRPK1 Inhibitors as Potential Topical Therapeutics for Neovascular Eye Disease. ACS Chem. Biol. 12, 825–832 (2017).

41. J. M. Hatcher et al., SRPKIN-1: A Covalent SRPK1/2 Inhibitor that Potently Converts VEGF from Pro-angiogenic to Anti-angiogenic Isoform. Cell chemical biology 25, 460–470.e466 (2018).

42. P. Wang et al., Both decreased and increased SRPK1 levels promote cancer by interfering with PHLPP-mediated dephosphorylation of Akt. Mol. Cell 54, 378–391 (2014).

43. F. Lamers et al., Knockdown of survivin (BIRC5) causes apoptosis in neuroblastoma via mitotic catastrophe. Endocr. Relat. Cancer 18, 657–668 (2011).

44. S. B. McMahon, MYC and the control of apoptosis. Cold Spring Harb. Perspect. Med. 4, a014407–a014407 (2014).

45. J. Campbell et al., Large-Scale Profiling of Kinase Dependencies in Cancer Cell Lines. Cell Reports 14, 2490–2501 (2016).

46. M. Bantscheff et al., Quantitative chemical proteomics reveals mechanisms of action of clinical ABL kinase inhibitors. Nat. Biotechnol. 25, 1035 (2007).

47. N. A. Karp et al., Addressing accuracy and precision issues in iTRAQ quantitation. Molecular & cellular proteomics: MCP 9, 1885–1897 (2010).

48. L. Ting, R. Rad, S. P. Gygi, W. Haas, MS3 eliminates ratio distortion in isobaric multiplexed quantitative proteomics. Nat Methods 8, 937–940 (2011).

49. A. Hogrebe et al., Benchmarking common quantification strategies for large-scale phosphoproteomics. Nature communications 9, 1045–1045 (2018).

50. M. M. Savitski et al., Multiplexed Proteome Dynamics Profiling Reveals Mechanisms Controlling Protein Homeostasis. Cell 173, 260–274.e225 (2018).

51. H. P. Gunawardena et al., QuantFusion: Novel Unified Methodology for Enhanced Coverage and Precision in Quantifying Global Proteomic Changes in Whole Tissues. Molecular & cellular proteomics: MCP 15, 740–751 (2016).

52. J. Cox et al., Accurate Proteome-wide Label-free Quantification by Delayed Normalization and Maximal Peptide Ratio Extraction, Termed MaxLFQ. Molecular & Cellular Proteomics: MCP 13, 2513–2526 (2014).

53. J. L. Hecht, G. L. Mutter, Molecular and pathologic aspects of endometrial carcinogenesis. J Clin Oncol 24, 4783–4791 (2006).

54. E. Kuhn et al., Identification of molecular pathway aberrations in uterine serous carcinoma by genome-wide analyses. J Natl Cancer Inst 104, 1503–1513 (2012).

55. K. Tzelepis et al., SRPK1 maintains acute myeloid leukemia through effects on isoform usage of epigenetic regulators including BRD4. Nature communications 9, 5378–5378 (2018).

56. E. Bonazzoli et al., Inhibition of BET Bromodomain Proteins with GS-5829 and GS-626510 in Uterine Serous Carcinoma, a Biologically Aggressive Variant of Endometrial Cancer. Clinical cancer research: an official journal of the American Association for Cancer Research 24, 4845–4853 (2018).

57. C. Di Tucci et al., Immunotherapy in endometrial cancer: new scenarios on the horizon. J. Gynecol. Oncol. 30, e46–e46 (2019).

58. P. A. Ott et al., Safety and Antitumor Activity of Pembrolizumab in Advanced Programmed Death Ligand 1-Positive Endometrial Cancer: Results From the KEYNOTE-028 Study. Journal of clinical oncology: official journal of the American Society of Clinical Oncology 35, 2535–2541 (2017).

59. L. Frankiw, D. Baltimore, G. Li, Alternative mRNA splicing in cancer immunotherapy. Nature Reviews Immunology 19, 675–687 (2019).

