## Supplemental figures and legends for "Kinome Profiling of Primary Endometrial Tumors Using Multiplexed Inhibitor Beads and Mass Spectrometry Identifies SRPK1 As Candidate Therapeutic Target"

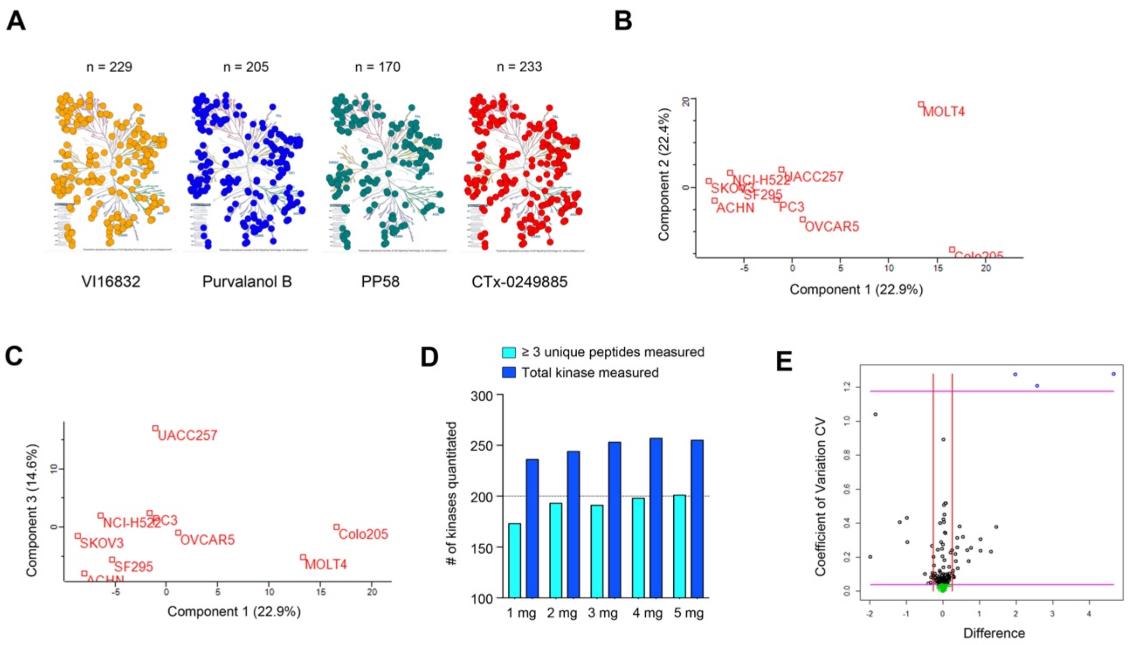


**Supplemental Figure 1. Designing a super-SILAC kinase standard for measuring kinase abundance in tissues.** A, Kinome trees depict the fraction of kinome enriched by each inhibitor resin. A mixture of cancer cell lines (5-mix used in for the SKS) was applied to each inhibitor resin and captured kinases detected by MIB-MS. B-C, Principle component analysis of SILAC-determined kinase log2 ratios of selected cancer cell lines. Each SILAC-labeled cell line was mixed 1:1 with the pooled non-labeled samples and analyzed by MIB-MS. D, Bar graph depicts the number of kinases detected from various protein concentration inputs. Various protein concentration of non-labeled OVCAR5 cells were mixed 1:1 with SKS and kinome(s) measured by MIB-MS. E, Reproducibility of the Q-MIBs assay was confirmed using a model-based approach for assessing technical reproducibility and outlier detection. A scatter plot of the coefficient of variation (CV) versus difference (D) for 300 duplicate MIB pairs for a particular Q-MIBs sample, demonstrates the reproducibility of the assay.

**A**

| Sample ID | Sample Type | Tumor Type | Protein input | # kinases measured | Filter out <180 kinases |
|---|---|---|---|---|---|
| N1 | Normal Endometrium | Normal | 5 mg | 223 | used in tumor vs normal |
| N2 | Normal Endometrium | Normal | 5 mg | 236 | used in tumor vs normal |
| N3 | Normal Endometrium | Normal | 5 mg | 253 | used in tumor vs normal |
| N4 | Normal Endometrium | Normal | 5 mg | 247 | used in tumor vs normal |
| N5 | Normal Endometrium | Normal | 5 mg | 234 | used in tumor vs normal |
| N6 | Normal Endometrium | Normal | 5 mg | 231 | used in tumor vs normal |
| N7 | Normal Endometrium | Normal | 5 mg | 227 | used in tumor vs normal |
| N8 | Normal Endometrium | Normal | 5 mg | 237 | used in tumor vs normal |
| N9 | Normal Endometrium | Normal | 5 mg | 248 | used in tumor vs normal |
| N10 | Normal Endometrium | Normal | 5 mg | 242 | used in tumor vs normal |
| N11 | Normal Endometrium | Normal | 5 mg | 224 | used in tumor vs normal |
| N12 | Normal Endometrium | Normal | 6 mg | N/A | insufficient kinase recovery |
| N13 | Normal Endometrium | Normal | 5 mg | 135 | filter out <180 kinases |
| N14 | Normal Endometrium | Normal | 5 mg | 230 | used in tumor vs normal |
| N15 | Normal Endometrium | Normal | 5 mg | 159 | filter out <180 kinases |
| N16 | Normal Endometrium | Normal | 5 mg | N/A | insufficient kinase recovery |
| N17 | Normal Endometrium | Normal | 5 mg | N/A | insufficient kinase recovery |
| N18 | Normal Endometrium | Normal | 5 mg | 228 | used in tumor vs normal |
| N19 | Normal Endometrium | Normal | 5 mg | N/A | insufficient kinase recovery |
| N20 | Normal Endometrium | Normal | 5 mg | 236 | used in tumor vs normal |
| T1 | Endometrial Tumor | Endometrioid | 5 mg | 236 | used in tumor vs normal |
| T2 | Endometrial Tumor | Endometrioid | 5 mg | 253 | used in tumor vs normal |
| T3 | Endometrial Tumor | Endometrioid | 5 mg | 237 | used in tumor vs normal |
| T4 | Endometrial Tumor | Endometrioid | 5 mg | 248 | used in tumor vs normal |
| T5 | Endometrial Tumor | Serous | 5 mg | 228 | used in tumor vs normal |
| T6 | Endometrial Tumor | Endometrioid | 5 mg | 247 | used in tumor vs normal |
| T7 | Endometrial Tumor | Endometrioid | 5 mg | 225 | used in tumor vs normal |
| T8 | Endometrial Tumor | Endometrioid | 5 mg | 241 | used in tumor vs normal |
| T9 | Endometrial Tumor | Endometrioid | 5 mg | 105 | filter out <180 kinases |
| T10 | Endometrial Tumor | Endometrioid | 5 mg | 75 | filter out <180 kinases |
| T11 | Endometrial Tumor | Endometrioid | 5 mg | 153 | filter out <180 kinases |
| T12 | Endometrial Tumor | Endometrioid | 5 mg | 206 | used in tumor vs normal |
| T13 | Endometrial Tumor | Endometrioid | 5 mg | 212 | used in tumor vs normal |
| T14 | Endometrial Tumor | Serous | 5 mg | 198 | used in tumor vs normal |
| T15 | Endometrial Tumor | Serous | 5 mg | 238 | used in tumor vs normal |
| T16 | Endometrial Tumor | Endometrioid | 5 mg | 242 | used in tumor vs normal |
| T17 | Endometrial Tumor | Endometrioid | 5 mg | 252 | used in tumor vs normal |
| T18 | Endometrial Tumor | Endometrioid | 5 mg | 243 | used in tumor vs normal |
| T19 | Endometrial Tumor | Endometrioid | 5 mg | 226 | used in tumor vs normal |
| T20 | Endometrial Tumor | Endometrioid | 5 mg | 216 | used in tumor vs normal |

**B**

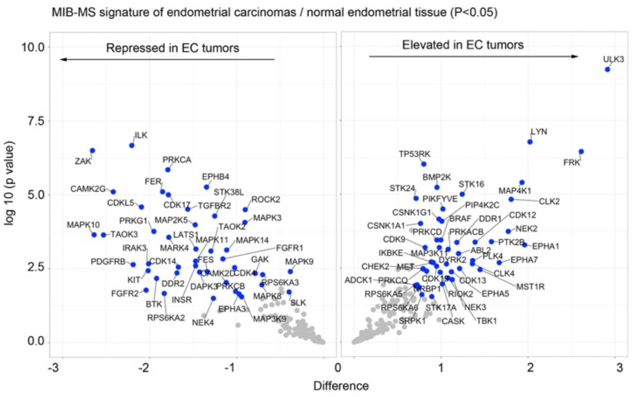


**Supplemental Figure 2. Kinome profiling of EC tumors and NE tissues using Q-MIBs.** A, Table depicts characteristics of EC tumors and NE tissues used in the kinome profiling experiments depicted in Figure 4. B, Volcano plot depicts kinases elevated or reduced in EC tumors relative to NE tissues. Statistical differences in kinase log2 s-SILAC ratios comparing EC tumors relative to NE tissues were determined by paired t-test P <0.05. Kinase log2 s-SILAC ratios were determined by comparing ratio of ratios (EC tumors /s-SILAC relative NE tissues /s-SILAC). MIB-MS profiling was performed in biological duplicate.

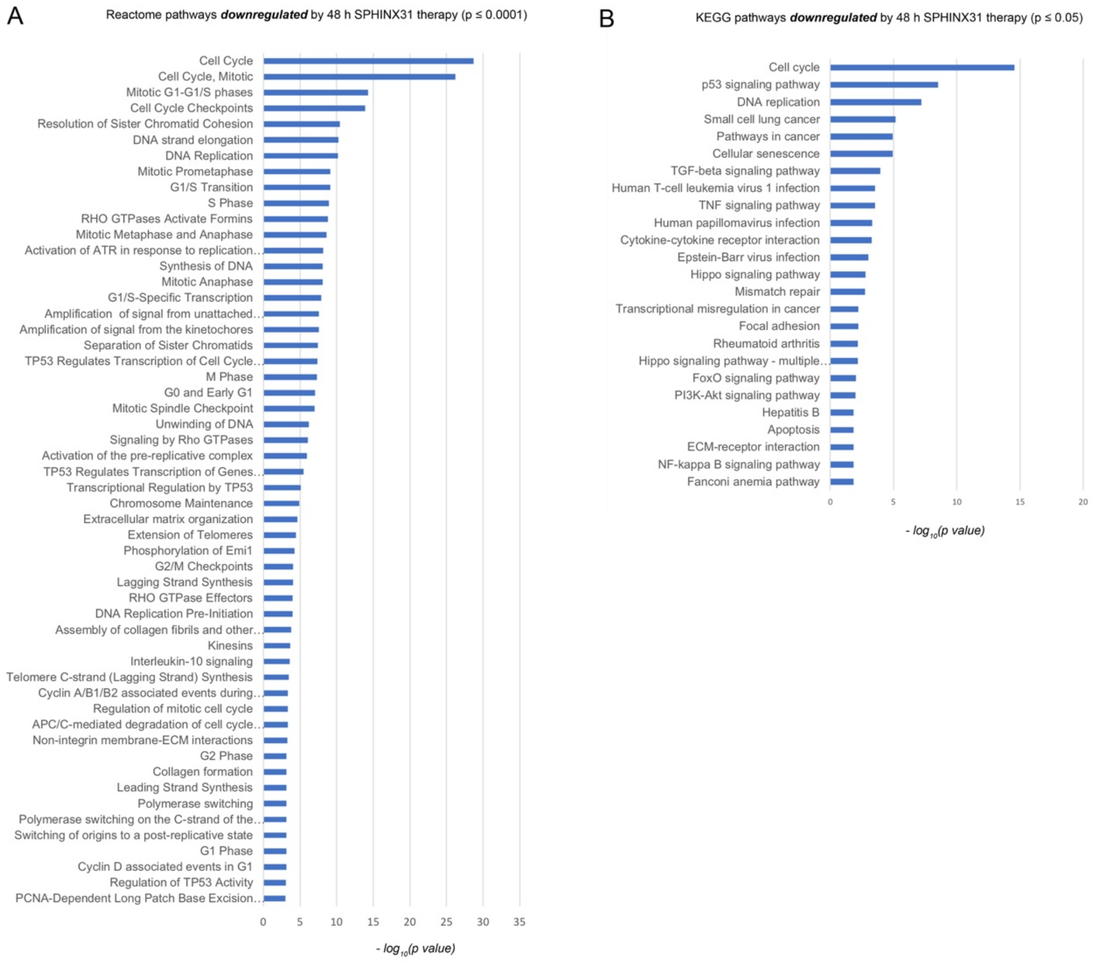


**Supplemental Figure 3. Pathway analysis of genes downregulated by SRPK1 inhibition in USC cells**. A-B, Genes significantly repressed by 24 h SPHINX31 (5 μM) treatment were analyzed by gProfiler to determine pathway enrichments (BH P<05). Bar plot depicts Reactome (A) and KEGG (B) pathways repressed by 48 h SPHINX31 therapy.
